# NUFEB: A Massively Parallel Simulator for Individual-based Modelling of Microbial Communities

**DOI:** 10.1101/648204

**Authors:** Bowen Li, Denis Taniguchi, Jayathilake Pahala Gedara, Valentina Gogulancea, Rebeca Gonzalez-Cabaleiro, Jinju Chen, Andrew Stephen McGough, Irina Dana Ofiteru, Thomas P Curtis, Paolo Zuliani

## Abstract

We present NUFEB, a flexible, efficient, and open source software for simulating the 3D dynamics of microbial communities. The tool is based on the Individual-based Modelling (IbM) approach, where microbes are represented as discrete units and their behaviour changes over time due to a variety of processes. This approach allows us to study population behaviours that emerge from the interaction between individuals and their environment. NUFEB is built on top of the classical molecular dynamics simulator LAMMPS, which we extended with IbM features. A wide range of biological, physical and chemical processes are implemented to explicitly model microbial systems. NUFEB is fully parallelised and allows for the simulation of large numbers of microbes (10^7^ individuals and beyond). The parallelisation is based on a domain decomposition scheme that divides the domain into multiple sub-domains which are distributed to different processors. NUFEB also offers a collection of post-processing routines for the visualisation and analysis of simulation output. In this article, we give an overview of NUFEB’s functionalities and implementation details. We provide examples that illustrate the type of microbial systems NUFEB can be used to model and simulate.

**Author summary:** Individual-based Models (IbM) are one of the most promising frameworks to study microbial communities, as they can explicitly describe the behaviour of each cell. The development of a general-purpose IbM solver should focus on efficiency and flexibility due to the unique characteristics of microbial systems. However, available tools for these purposes present significant limitations. Most of them only facilitate serial computing for single simulation, or only focus on biological processes, but do not model mechanical and chemical processes in detail. In this work, we introduce the IbM solver NUFEB that addresses these shortcoming. The tool facilitates the modelling of much needed biological, chemical, physical and individual microbes in detail, and offers the flexibility of model extension and customisation. NUFEB is also fully parallelised and allows for the simulation of large complex microbial system. In this paper, we first give an overview of NUFEB’s functionalities and implementation details. Then, we use NUFEB to model and simulate a biofilm system with fluid dynamics, and a large and complex biofilm system with multiple microbial functional groups and multiple nutrients.

This is a *PLOS Computational Biology* Software paper.

## Introduction

Microbial communities are groups of microbes that live together in a contiguous environment and interact with each other. The presence of microbial communities on the planet plays an important role in natural processes, as well as in environmental engineering applications such as wastewater treatment [1], waste recycling [2] and the production of alternative energy [3]. Therefore, studies on how these communities form and evolve have become increasingly important over the past few decades [4,5].

Work on microbial communities has revealed that the community’s emergent behaviour arises from a variety of interactions between microbes and their local environment. *In vitro* experiments offer a way to gain insights into these complex interactions, but at great expense in time and resources. On the other hand, *in silico* computational models and numerical simulations could help researchers to investigate and predict how complex processes affect the behaviour of biological systems in an explicit and efficient way. Different approaches have been developed for modelling microbial communities [6–8]. One of the most promising strategies is to develop a mathematical model from the description of the characteristics of the individual microbes, usually referred to as Individual-based Models (IbM) [9,10]. In conventional IbM, the microbes are represented as rigid particles, each of which is associated with a set of properties such as mass, position, and velocity. These properties are affected by internal or external processes (e.g., diffusion), resulting in microbial growth, decay, motility, etc. Therefore, IbM are particularly useful when one is interested in understanding how individual heterogeneity and local interactions influence an emergent behaviour.

The development of a general-purpose IbM solver should focus on the following aspects. First, the solver needs to be flexible. Depending on the purpose of the model, IbM may involve multiple microbial functional groups, nutrients and sophisticated biological, chemical and physical processes, or sometimes it may be a simple model that describes mono-functional group or focuses on a few processes. Thus, it is important for the solver to be highly customisable (for building IbM) and extendible (with new IbM features). Second, the solver should be scalable. Simulation of large microbial communities is difficult since they contain a very high number of individuals. Different modelling strategies have been proposed to overcome this limitation, including using super-individuals and statistically representative volume elements [11]. However, there is very little work on the development of a scalable IbM solver. Parallel computing can help scalability by using multiple computer resources to simulate many individuals simultaneously. This is accomplished by breaking the problem domain into discrete regions which separate out the individuals as much as possible, allowing each processing element to simulate the local interactions between individuals whilst minimizing the interactions between regions. In this way each region can largely run concurrently with the others.

In this work, we present a three-dimensional, open-source, and massively parallel IbM solver called NUFEB that addresses these desired above. The purpose of NUFEB is to offer a flexible and efficient framework for simulating microbial communities at the micro-scale. A comprehensive IbM is implemented in the solver which explicitly models biological, chemical and physical processes, as well as individual microbes. The present solver supports parallel computing and allows flexible extension and customisation of the model. NUFEB is based on the state-of-the-art software LAMMPS (Large-scale Atomic Molecular Massively Parallel Simulator) [12]. We selected LAMMPS because of its open-source, parallel, and extendible nature. There are several open-source IbM solvers that have been developed over the past decade and widely applied to microbiology research, such as iDynoMiCS [13], SimBiotics [14], BioDynaMo [15], and DiSCUS [16]. However, most of them only facilitate serial computing for single simulation, or focus only on biological processes, but do not model mechanical and chemical processes in detail (e.g., fluid dynamics, mechanical interaction, pH dynamics, and thermodynamics). The NUFEB simulator instead includes all of these features.

## Model description

In this section, we describe and review the IbM implemented in NUFEB. We build on the ideas described previously in [17] and extend them to cope with hydrodynamics and chemical processes. For the sake of accuracy, in the following we use the term “microbe” when describing biological and chemical processes, and use the term “particle” when describing physical process. However, the two terms effectively mean the same thing in the model.

### Computational domain

The *computational domain* is the environment where microbes reside and the biological, physical and chemical processes take place. It is defined as a micro-scale 3D rectangular box with dimensions *L_X_* × *L_Y_* × *L_Z_*. The size of the domain normally ranges from hundreds to thousands micrometers for micro-scale simulation. Therefore, the computational domain is considered as a sub-space of a macro-scale bioreactor or any other large-scale microbiological system. We assume that the macro-scale ecosystem is made up of replicates of the micro-scale domain if the bioreactor is perfectly mixed [17,18]. Within the domain, chemical properties such as nutrient concentration, pH and Gibbs free energy are represented as continuous fields. To resolve their dynamics over time and space, the domain is discretised into Cartesian grid elements so that the values can be calculated at each discrete voxel on the meshed geometry. Domain boundary conditions can be defined as either periodic or fixed. The former allows particles to cross the boundary, and re-appear on the opposite side of the domain, while a fixed wall prevents particles to interact across the boundary.

NUFEB allows the modelling and simulation of a biofilm system where different compartments may be defined within the computational domain [19] (Fig 1). The biofilm compartment is the volume occupied by microbial agents and their extracellular polymeric substances (EPS), in which the distribution of soluble chemical species is affected by both diffusion and reaction processes. The boundary layer compartment lies over the biofilm, so that nutrient diffusion and advection is resolved in this space. The bulk liquid compartment is situated at the top of the boundary layer, and nutrients in this area are assumed to be perfectly mixed with the same concentration as in the macro-scale bioreactor. We also assume that the boundary layer compartment stretches from the maximum biofilm thickness to the bulk liquid, with the height of the compartment specified by the user. The boundary between the diffusion layer and bulk liquid compartments is parallel with the bottom surface (substratum), but its location needs to be updated when the biofilm thickness changes.

**Fig 1.**
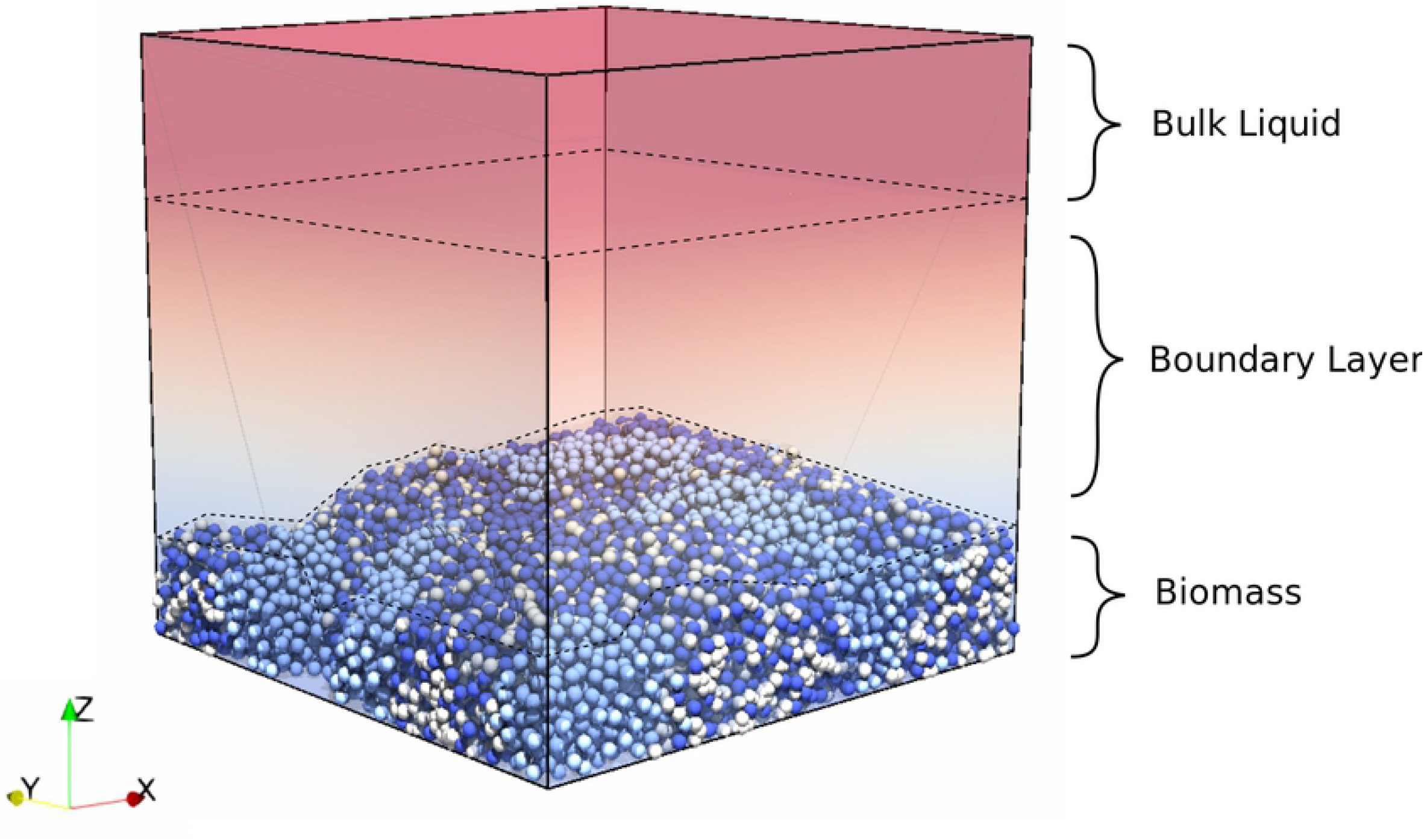
The computational domain with different compartments. The top region is the well mixed bulk liquid compartment, the middle region is the diffusion boundary layer compartment, and the bottom region represents the biofilm compartment.

### Biological processes

In the IbM, microbes are modelled as rigid spheres, with each individual having a set of state variables including position, density, velocity, force, functional group, mass, diameter, outer-mass and outer-diameter. These attributes vary among individuals and can change through time. Outer-diameter and outer-mass are used to represent an EPS shell: in some microbes EPS is initially accumulated as an extra shell around the particle. Microbial functional groups (types) are groups of one or more individual microbes that have the same or similar biological behaviour. The separation of individuals into different functional groups is based on their specific metabolism.

The NUFEB biological sub-model handles microbial metabolism, growth, decay, and reproduction (cell division and EPS formation). Details of the biological sub-model are given below.

#### Microbe growth and decay

An individual microbe grows and its mass increases by consuming nutrients supplied by the bulk liquid. The process of growth and decay is described by the following ordinary differential equation:

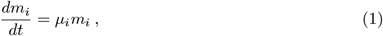

where *m_i_* is the biomass of the *i^th^* microbe, and *μ_i_* is the specified growth rate. To determine *μ_i_*, two growth models are implemented: (i) *Monod-based* and (ii) *energy-based.* The user can choose one of the growth models when configuring the simulation to run. For exemplification, the Monod-based growth model implements the work described in [17] and [18]. Three functional groups of microbes and two inert states are considered. They include: active heterotrophs (HET), ammonia oxidizing bacteria (AOB), nitrite oxidizing bacteria (NOB), and inactive EPS and dead cells. Microbial growth is based on Monod kinetics driven by the local concentration of nutrients (*S*_substrate_, *S*_NH_4_+_, *S*_NO_2_-_, *S*_O_2__) at the voxel in which each microbe resides [20]. The decay rate is assumed to be first order.

The *energy-based growth model* implements the work proposed in [21]. In this model, the growth rate of each microbe *μ_i_* is determined not only by nutrient availability but also by the amount of energy available for its metabolism:

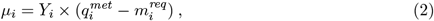

where the maximum growth yield *Y_i_* is estimated by using the Energy Dissipation Method [22], 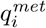 is the metabolic rate which depends on the availability of nutrients (in particular, their dissociation forms), and 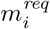 is the average maintenance requirement. Thus, a microbe grows if it harvests more energy than the necessary for its maintenance requirement. On the other hand, the microbe decays when the energy requirement is not met.

#### Microbe division and death

Microbe division is the result of biomass growth, while death is the result of biomass decay. Both are considered as instantaneous processes. Division occurs if the diameter of a microbe reaches a user-specified threshold value; the cell then divides into two daughter cells. The total mass of the two daughter cells is always conserved from the parent cell. One daughter cell is (uniformly) randomly assigned 40% – 60% of the parent cell’s mass, and the other gets the rest. Also, one daughter cell takes the location of the parent cell while the centre of the other daughter cell is (uniformly) randomly chosen at a distance *d* (distance between the centres of the two agents) corresponding to the sum of the diameter of the daughters.

The size of a microbe decreases when nutrients are limited or the energy available is not sufficient to meet maintenance requirements. Microbes which shrink below a user-specified minimum diameter are considered as dead. Their type then changes to the dead type. Dead cells do not perform biological activities but their biomass will linearly convert to carbon. For computational efficiency, decaying cells are removed from the system when their size is sufficiently small (defined as 10 times smaller than the death threshold diameter).

#### EPS production

The Monod-based growth model allows active heterotrophs to secrete EPS into their neighbouring environment. The EPS play an important role in microbial aggregation by offering a protective medium. The production process follows the approach presented in [17] and [23] with the simplification that EPS are secreted by heterotrophs only. Initially, EPS is accumulated as an extra shell around a HET particle (note that EPS density is lower than microbe density). When the relative thickness of the EPS shell of the HET particle exceeds a certain threshold value, almost half (uniformly random ratio between 0.4-0.6) of the EPS mass excretes as a separate EPS particle and is (uniformly) randomly placed next to the HET.

### Physical processes

NUFEB’s physical sub-model includes two key features and the dependencies between them: microbes (particles) and fluid. Microbes interact among themselves and with the ambient fluid. The physics of microbial motion is solved by using the discrete element method (DEM). Fluid momentum and continuity equations are solved based on computational fluid dynamics (CFD) and coupled with particle motion.

#### Mechanical relaxation

When microbes grow and divide, the system may deviate from mechanical equilibrium (i.e., non-zero net force on particles) due to particle overlap or collision. Hence, mechanical relaxation is required to update the location of the particles and minimise the stored mechanical energy of the system. Mechanical relaxation is carried out using the discrete element method, and the Newtonian equations of motion are solved for each particle in a Lagrangian framework. The equation for the translational and rotational movement of particle *i* is given by:

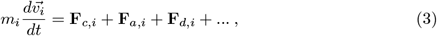

where *m_i_* is the mass, and 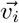 is the velocity. The type of force acting on the particle varies according to different biological systems. For example, the above equation takes into account three commonly used forces in microbial system. The contact force **F**_*c,i*_ is a pair-wise force exerted on the particles to resolve the overlap problem at the particle level. The force equation is solved based on Hooke’s law, as described in [24]:

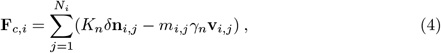

where *N_i_* is the total number of neighbouring particles of *i, K_n_* is the elastic constant for normal contact, *δ***n***_i,j_* are overlap distance between the center of particles *i* and its neighbour particle *j, m_i,j_* is the effective mass of particles *i* and *j*, *γ_n_* is the viscoelastic damping constant for normal contact, and **v**_*i,j*_ is the relative velocity of the two particles.

The EPS adhesive force **F**_*a,i*_ is a pair-wise interaction, which is modelled as a van der Waals force [25]:

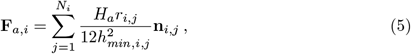

where *H_a_* is the Hamaker coefficient, *r_i,j_* is the effective radius of particles *i* and *j*, *h_min,i,j_* is the minimum separation distance of the two particles, and **n**_*i,j*_ is the unit vector from particle *i* to *j*.

The drag force **F***_d,i_* is the fluid-particle interaction force due to fluid flow, with direction opposite the microbe motion in a fluid. It is formulated as [25]:

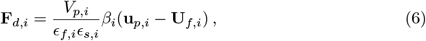

where *ϵ_s,i_* is the particle volume fraction, *ϵ_f,i_* = 1 − *ϵ_s,i_* is the fluid volume fraction, *V_p,i_* and **u**_*p,i*_ are volume and velocity of particle *i*, respectively, **U**_*f,i*_ is the fluid velocity imposed on particle *i*, and *β_i_* is the drag correction coefficient [26]. Apart from the forces above, the LAMMPS framework offers mechanical interactions that one can apply directly to IbM (see LAMMPS’ user manual for more details [27]).

#### Fluid dynamics

Hydrodynamics is an important factor in microbial community modelling as microbes usually live in water where their behaviour is influenced by fluid flow in two ways: transport of nutrients and detachment. Nevertheless, accurate hydrodynamics has rarely been considered in 3D microbial community modelling due to its computational complexity [28]. In NUFEB, with the support of code parallelisation, hydrodynamics is introduced by using the CFD-DEM approach [29,30]. In this approach, DEM solves the motion of Lagrangian particles based on Newton’s second law, while CFD (computational fluid dynamics) tracks the motion of fluid based on locally averaged Navier-Stokes equations.

The fluid velocity at each voxel in space is replaced by its average, and the locally averaged incompressible continuity and momentum equations for the fluid phase are given by [31]:

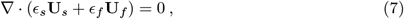

and

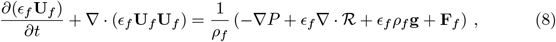

where *ϵ_s_*, **U**_*s*_, and **F**_*f*_ are the fields of the solid volume fraction, velocity and fluid-particle interaction forces (e.g., drag force) of microbes, respectively. They are obtained by averaging discrete particle data in DEM [32,33]; *ϵ_f_* is the fluid volume fraction and **U**_*f*_ is the fluid velocity. Besides the fluid-particle interaction, the terms on the right-hand side of Eq 8 also include the fluid density *ρ_f_*, the pressure gradient ∇*P*, the divergence of the stress tensor 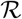 and the gravitational acceleration **g**. The fluid momentum equations are discretised and solved on an Eulerian grid by a finite volume method. The results, in particular the velocity field and the particle drag force, are used for solving nutrient transport and mechanical relaxation, respectively.

### Chemical processes

Nutrient transport is described using the diffusion-advection-reaction equation. To improve the representative of microbial growth, NUFEB also allows pH dynamics and gas-liquid transfer to be considered. In this section, we briefly address the main ideas of this chemical sub-model.

#### Nutrient consumption

The rate of nutrient consumption (or reaction rate) is calculated at each voxel. The reaction rates in the Monod-based growth model are defined according to Monod kinetics, where the stoichiometric matrix for particulate and soluble components is given in the S1 File. By contrast, in the energy-based growth model the consumption/formation rate of the *i^th^* microbe for each soluble species is driven by the microbial growth rate and the stoichiometric coefficients of the overall growth reaction, and is formalised as follows [21]:

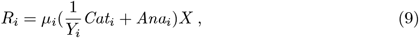

where *Cat_i_* is the free energy supplied by the microbial catabolic reactions, *Ana_i_* is the free energy required by the anabolic reactions, and *X* is the biomass density.

#### Nutrient mass balance

Nutrient concentration at each point within the computational domain is affected by different processes. To solve the nutrient distribution for each soluble component, the following diffusion-advection-reaction equation for the solute concentration *S* is employed in the model:

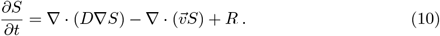

On the right-hand side of the equation, the first part is the diffusion term which describes nutrient movement from a region of high concentration to a region of low concentration, where ∇ is the gradient operator and *D* is the diffusion coefficient. The second part is the advection term which describes nutrient motion along the fluid flow, where 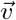 is the fluid velocity field. Finally, *R* is the reaction term which is governed by both biological activities (e.g., microbial growth causing nutrient consumption) and chemical activities (e.g., solute component transferred into gas).

Nutrient concentration in the bulk compartment is dynamically updated according to the following mass balance equation [19]:

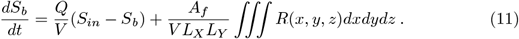

The bulk concentration *S_b_* of each soluble component is influenced by the nutrient inflow and outflow in the bioreactor (*Q* is the volumetric flow rate, *V* is the bioreactor volume and *S_in_* is the influent nutrient concentration), as well as the total consumption rate in the biofilm volume in the bioreactor (*A_f_* is the biofilm surface area, and *R*(*x, y, z*) is the reaction rate at each voxel.).

#### pH dynamics

We assume the influent enters the bulk liquid with a fixed pH value. However, the pH varies in space and time due to change in nutrient concentration as a result of microbial activity. The model considers acid-base reactions as equilibrium processes. The kinetic expressions are amended to consider only the non-charged form of the nutrients (e.g., HNO_2_ but not NO_2_^−^). The concentration of all dissociated and undissociated forms can be expressed as a function of the proton (H^+^) concentration. Then, the proton concentration can be determined by finding the root between 1 and 10^−14^ of the following charge balance equation, using an implicit Newton-Raphson approximation:

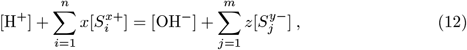

where *n* and *m* are the total number of cations and anions contributing to the pH, respectively, *x* and *y* are the charges corresponding to the cations and anions considered in the dissociation equilibrium, and *S* is the concentrations of cations and anions calculated based on the Gibbs free energy and temperature.

#### Gas-liquid transfer

A gas field can be defined in NUFEB to describe the rate of nutrient mass transfer from gas to liquid or vice-versa. The equilibrium between liquid and gas is disturbed by the acid-base reaction and the microbial activity taking place in the biofilm and bulk liquid. On the other hand, mass transfer may also affect microbial growth due to varying nutrient concentration in the liquid phase. The reaction rate from gas to liquid *R_G→L_* of a given nutrient can be expressed by the following equation [34]:

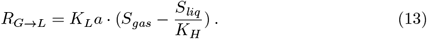

The rate is determined by the mass transfer coefficient of chemical component *K_L_a*, the gas concentration *S_gas_* in the head space, the saturation liquid concentration *S_liq_*, and Henry’s constant *K_H_*. Mass transfer from liquid to gas is formalised as follows:

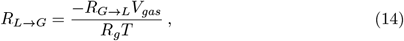

where *V_gas_* is the volume of the reactor head space, which is considered of equal size to the computational domain, and *R_g_* and *T* are the ideal gas constant and temperature, respectively.

### Design and implementations

NUFEB is developed in C++ as a user package within the LAMMPS platform. A prototype implementation was used in [17] to study physical behaviour of microbial communities. We describe a major improvement of the tool, in which new features and enhancements have been developed including coupling with fluid dynamics, chemical processes, code parallelisation, and post-processing routines. In this section, we summarise the NUFEB functionalities and give some implementation details.

### IbM in LAMMPS

NUFEB is built on top of LAMMPS and extends it with IbM features. LAMMPS is a classical molecular dynamics simulator and primarily solves particle physics, including a wide range of inter-particle interactions and potentials [12]. In NUFEB, a new sphere-like particle type is defined with additional attributes (e.g., outer-diameter, outer-mass) to represent a microbe. Microbes are grouped into different functional groups, and members of each group share the same biological parameters. The computational domain is restricted to a 3D rectangular box with user-specified dimensions, Cartesian grid size and boundary conditions. In LAMMPS, a “fix” is any operation that applies to the system during time integration. Examples include the updating of particle locations, velocities, forces. NUFEB defines a series of new fixes to perform the IbM related processes previously described. The user can also easily extend the tool with other processes by adding new fix commands. During the simulation, fixes are invoked at a user-defined frequency to update field quantities and microbe attributes. For the purpose of computational efficiency and different modelling systems, NUFEB allows users to customise a simulation by enabling/disabling any of the fix commands from the input setting.

In order to execute a NUFEB simulation, an input script (a text file) is prepared with certain commands and parameters. NUFEB will read those commands and parameters, one line at a time. Each command causes NUFEB to take some action, such as setting initial conditions, performing IbM processes, or running a simulation. In Fig 2, we show an excerpt of an input script used in this work (Case Study 2). The details of the input format are described in S1 Manual.

**Fig 2.**
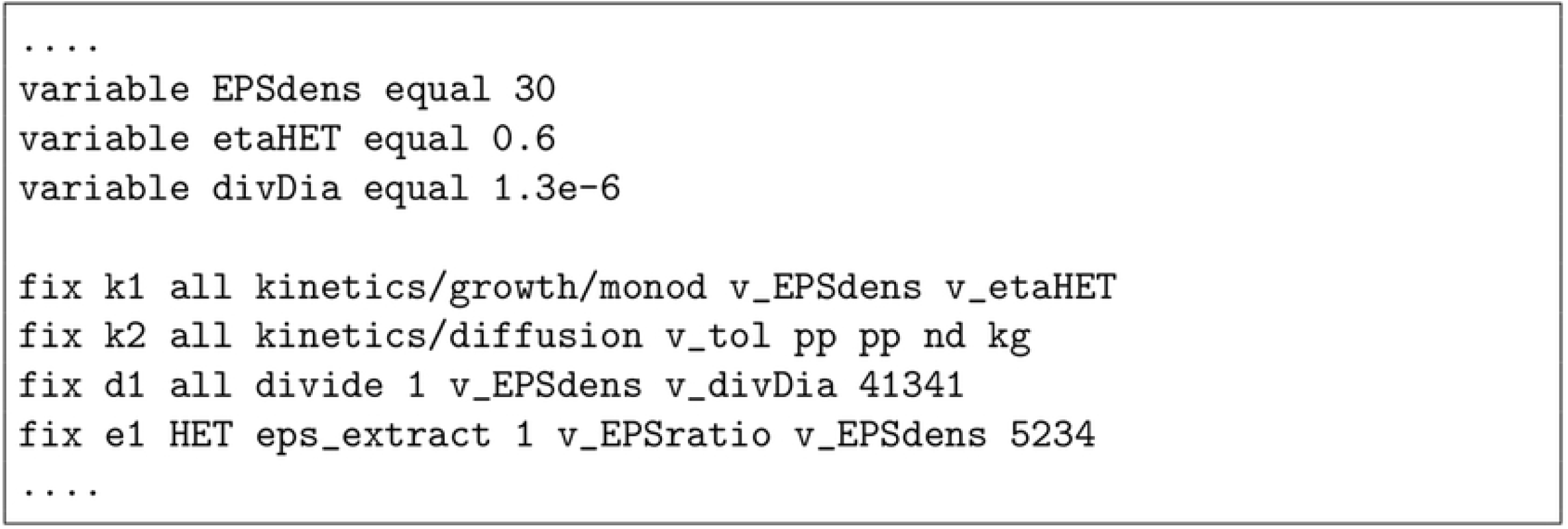
An example of (partial) input script for NUFEB simulation. A list of “fix” commands defines IbM processes that apply to the simulation, which includes Monod-based growth (`k1`), nutrient mass balance (`k2`), cell division (`d1`), and EPS production (`el`). Each fix command may require one or more parameters for the model specification, such as EPS density (`EPSdens`), HET reduction factor in anoxic condition (`etaHET`), and division diameter (`divDia`).

The procedure of a classical IbM simulation in NUFEB is presented in Algorithm 1. The model’s processes can be operated sequentially as they are on different timescales. The mechanical timestep is typically of the order of 10^−7^s; the diffusion timestep is of the order of 10^−4^s, while the biological timestep is much larger ranging from minutes to hours. The coupling between multiple timescales relies on the pseudo steady-state approximation and the frozen state [28]. For example, when a steady state solute concentration is reached at each biological timestep, the concentration is assumed to remain unchanged (frozen state) until the next biological step. In this way, the computational load for solving fast dynamic processes can be significantly reduced.

#### Algorithm 1 (IbM simulation procedure)

~~~
1: **input:** initial states of computation domain, microbes and fields
2: **output:** states of all microbes and fields at each output time step
3: **while** biological time step *t_bio_* < *t_end_* **do**
4:   solve fluid dynamics to update velocity field and particle (drag) force
5:   solve nutrient mass balance to update solute concentration field ^1^
6:   update nutrient concentration in bulk compartment
7:   perform microbe growth to update biomass and size
8:   perform microbe division, death and EPS production
9:   perform mechanical relaxation to update microbe position and velocity 10: update boundary layer location and neighbour list
11:  *t_bio_* = *t_bio_* + Δ*t*
12: **end while**
~~~

Mechanical relaxation is resolved by using the Verlet algorithm provided by LAMMPS [35]. The computation of interaction forces between particles, such as contact force, relies on LAMMPS’ neighbour lists. During the Verlet integration, instead of iterating through every other particle, which would result in a quadratic time complexity algorithm, LAMMPS maintains a list of neighbours for each particle. The computation of the interaction force is only performed between a particle and its corresponding neighbours. The neighbour lists must be updated from time to time depending on the microbe’s displacement and division.

The nutrient mass balance equation is discretised on a Marker-And-Cell (MAC) uniform grid and the concentration scalar is defined at the centre of the voxel (cubic grid element). The temporal and spatial derivatives of the transport equation are discretised by Forward Euler and Central Finite Differences, respectively. Depending on the physical situation, different boundary conditions can be chosen when solving the equation. For instance, a biofilm system would normally uses non-flux Neumann conditions through the bottom surface to model impermeable support material, Dirichlet boundary conditions at the top surface as it is assumed to connect with bulk environment, and periodic boundary conditions for the rest of the four surfaces [19].

### Coupling with fluid dynamics

NUFEB employs and extends the existing CFD-DEM solver SediFoam for the simulation of hydrodynamics [36]. SediFoam provides a flexible interface between the two open-source solvers LAMMPS and OpenFOAM. LAMMPS aims to simulate particle motions, while OpenFOAM (Open Field Operation and Manipulation) is a parallel CFD solver that can perform three-dimensional fluid flow simulations [37]. Inbetween them, SediFoam offers efficient parallel algorithms that transfers and maps the properties of Lagrangian particles to an Eulerian mesh, and vice versa. In this work, we also extend SediFoam for compatibility with IbM features, in particular, to transfer and map the information of new divided particles and velocity field between the solvers.

The schema of the CFD-DEM coupling is shown in Fig 3. At each fluid timestep, particle information (e.g., mass, force, velocity, tag) is transferred from the DEM module to the CFD module. A particles list maintained by OpenFOAM is updated based on the obtained information. An averaging procedure is then performed to convert the properties of discrete particles to the Eulerian CFD mesh. The fluid momentum equations are discretised and solved on the Eulerian mesh by the finite volume method. The PISO (Pressure-Implicit with Splitting of Operators) algorithm is followed to solve the equations [38]. In the DEM module, the solutions of drag force and velocity field obtained from OpenFOAM are assigned to each particle and corresponding mesh grid, respectively. The drag force will be taken into account in evolving the motion of the particles. The velocity field is used for solving the advection-diffusion-reaction equation (Eq 10).

**Fig 3.**
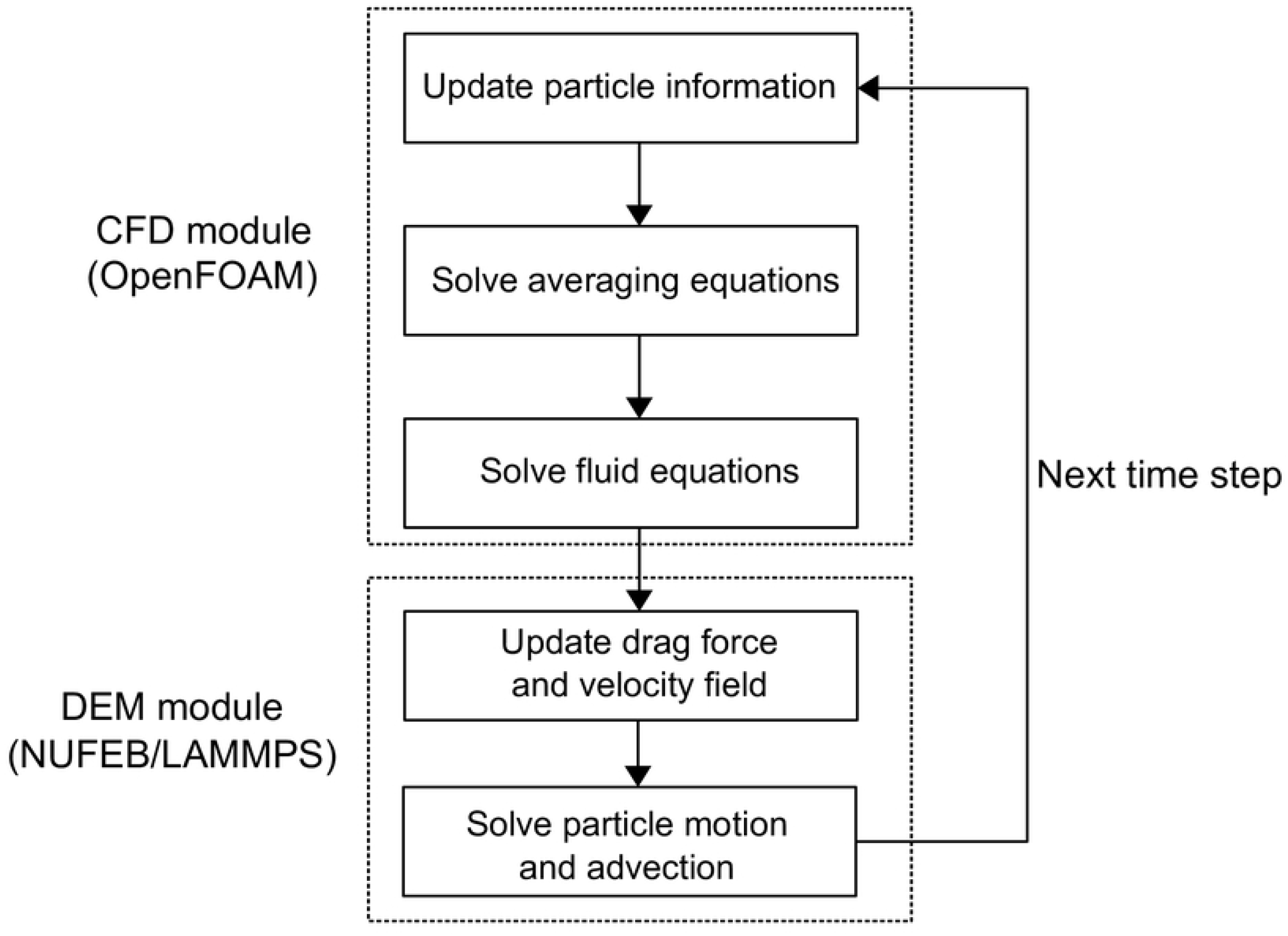
Block diagram of CFD-DEM coupling. Diagram adopted from [36]. Fluid dynamics is solved in the CFD module and particle motion is solved in the DEM module. Particle and field information are transferred between the two modules based on an averaging procedure.

### Code parallelisation

The IbM implementation in NUFEB involves particle-based functions (e.g., contact force, division, and EPS excretion) and continuum-based functions (e.g., nutrient mass balance, and pH). The parallelisation of the former function group is based on LAMMPS’ parallel mechanism. The computational domain is spatially decomposed into multiple MPI (Message Passing Interface) processes (sub-domains). Each sub-domain contains local particles as well as ghost particles. Local particles are those residing in the owned sub-domain, and each process is responsible for updating the status of their local particles. Ghost particles are copies of particles owned by neighbouring processes. During the simulation, the local particles obtain information from their ghost (and neighbour) counterparts for calculating and updating their physical properties (e.g., forces). The neighbour lists require updating when particles are deleted and created due to microbe decay, division, etc.

Continuum-based functions have their variables computed using a uniform grid. Parallelism is achieved by spatially decomposing the computational domain into sub-domains. Computations such as diffusion and advection require solute information from adjacent cells. Therefore, like particle-based functions, grid cells on the boundary of each sub-domain need to be communicated to neighbouring sub-domains and are treated locally as ghost cells. The implementation of grid decomposition also considers the sub-domain box to be always conforming to the uniform grid boundary.

Automatic vectorisation was employed to further optimise computation intensive routines, such as pH calculation. The loops in the routines are vectorised and can be performed simultaneously. The vectorisation is achieved together with the use of control directives (i.e., `#pragmas`) to instruct the compiler on how to handle data dependencies within a loop.

### Other features

NUFEB allows simulation results to be stored into various formats for visualisation or analysis. The supported formats are VTK, POVray and HDF5. The VTK binary format is readable by the VTK visualisation toolkit or other visualisation tools, such as ParaView (all biofilm figures shown in the Results section were produced from ParaView). A post-processing routine is implemented to convert the LAMMPS default format to POVray image, which supports high rendering of particles. HDF5 is a hierarchical, filesystem-like data format supported by a number of popular software platforms, including Java, MATLAB and Python. This allows the user to directly import simulation data to any of the platforms for further analysis.

To understand the morphological dynamics of microbial systems, characteristics such as biofilm average height, biofilm surface roughness, floc equivalent diameter and floc fractal dimension can be measured during simulation. These aggregated characteristics are essential factors to study and design microbial system [39]. Parallelisation of the characteristics measurements is supported by NUFEB and based on domain decomposition.

NUFEB also supports most of the LAMMPS default commands, which can be useful in microbial simulations. For example, the *restart* and *read restart* functions write out the current state of a simulation as a binary file, and then start a new simulation with the previously saved system configuration; the *lattice* and *create atoms* functions allow to automatically create large numbers of initial microbes based on a user-defined lattice structure; *load balance* is performed with the objective of maintaining the same number of particles in each sub-domain in parallel runs. During the simulation, this function adjusts the size and shape of the sub-domains to balance the computational cost in the processors.

## Results

We have successfully validated NUFEB against two biofilm benchmark problems BM2 and BM3 proposed by the International Water Association (IWA) task group on biofilm modelling [40,41]. The validation results can be found in the S1 File. This section shows two further examples implemented using NUFEB. The case studies are based on biofilm systems, which are microbial communities of single or multiple microbial functional groups in which cells stick to each other and attach to a substratum by means of EPS.

### Case Study 1: Biofilm deformation and detachment

One of the outstanding questions in biofilm research is understanding how fluid-biofilm interactions affect the mechanical properties of biofilms. In this section, we describe how to use NUFEB to simulate a biofilm system with fluid dynamics.

To simulate a hydrodynamic biofilm, we apply fluid flow to a pre-grown biofilm that consists of heterotrophs and their EPS production. The biofilm is grown from 40 microbes inoculated on the substratum to a pre-determined height (80 *μm*) without flow and in an oxygen-limited condition (1 × 10^−4^ kg m^−3^). In this way, a mushroom-shaped biofilm structure can be developed to model liquid filled voids and channels (see Fig 4(a) and S1 Video). Then, we impose a fluid flow to the biofilm. During the fluid stage, any biological process is considered to be in the frozen state due to the small time scale of hydrodynamic calculations, and nutrient mass balance is omitted for the sake of simplicity. The motion of microbes is driven by both particle-particle and particle-fluid interactions, including EPS adhesion, contact force and drag force, as described in Eqs (4)–(6). The physical model parameters are kept constant throughout the simulation and can be found in the S1 File. For boundary conditions, we impose a fixed velocity **U**_*f*_ at the top surface with direction along the x-axis as inlet velocity, no-slip condition at the bottom surface and periodic conditions at other four surfaces. The pressure are enforced as zero gradient at the top and bottom surfaces.

**Fig 4.**
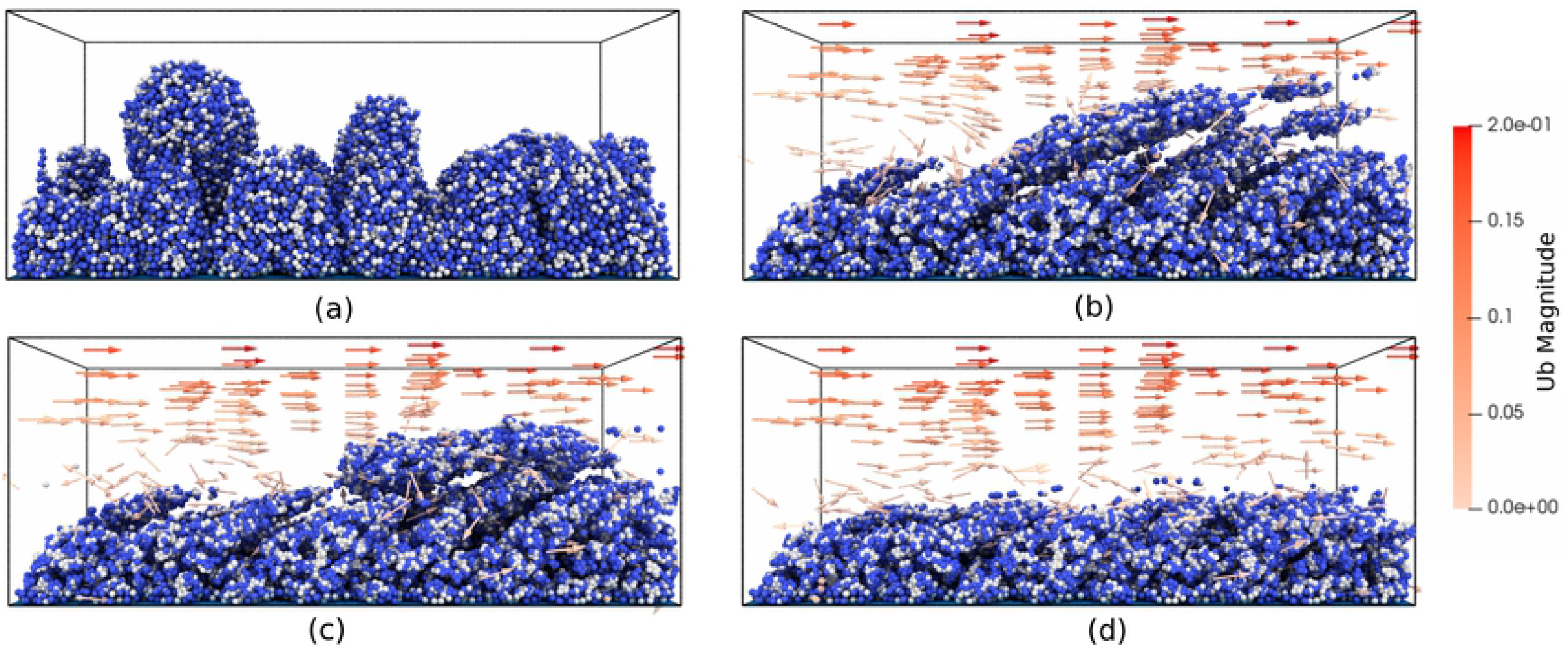
Biofilm deformation and detachment at U_*f*_ = 0.2 m s^−1^: (a) Time = 0; (b) Time = 0.0015s; (c) Time = 0.003s; (d) Time = 0.01s. The model simulates 4 × 10^4^ particles. Particles crossing the domain boundary will be removed from the system. Particle colours are blue for heterotrophs and grey for EPS.

Fig 4(b)-(d) and S2 Video show the biofilm deformation and detachment at **U**_*f*_ = 0.2 ms^−1^ (Reynolds number = 20). The biofilm deforms and microbes detach along with the flow direction. In the early stage of the detachment process, the top of the biofilm is highly elongated and forms filamentous streamers. However, most of the microbes are still connected together with cohesion, and there is only a small number of clusters detached from the head of the streamers due to cohesive failures (Fig 4(b)). As the fluid continues to flow, large chunks of microbes detach from the biofilm surface. These detached microbe chunks may also break-up again, re-agglomerate with other clusters or re-attach to the biofilm surface (Fig 4(c)). Such deformation and detachment events observed from our NUFEB simulation show qualitative agreement with both experimental results [42] and other numerical simulations using different methods [17,43]. The deformation reaches a pseudo-steady-state when the mushroom-shaped biofilm protrusions are removed from the system. As a result, the biofilm morphology changes dramatically from a rough to a flat surface (Fig 4(d)). During the deformation, the fluid, represented as red arrows, travels around the biofilm. Due to the irregular shape of the biofilm and the high fluid velocity, small vortexes can be observed at the biofilm surface on both the upstream and downstream sides. This phenomenon has been observed in previous studies [43].

For a more quantitative measurement of the deforming biofilm, we evaluated the total biomass and surface roughness of the biofilm at different fluid velocities. The biofilm surface roughness is calculated by [17]:

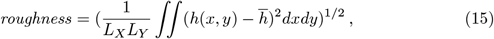

where *h*(*x, y*) is the biofilm height in the *z* direction at location (*x, y*) on the substratum, and 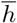 is the average biofilm height. As expected, when the fluid velocity increases the removed biomass also increases. For example, when **U**_*f*_ = 0.4 m s^−1^ is applied, the biomass reaches steady-state after 0.004s, and the total biomass decreases by 23%. By contrast, only 14% of the biomass can be removed if **U**_*f*_ = 0.2 m s^−1^ is applied (Fig 5(a)). The biofilm surface roughness shows a similar trend: the roughness decreases with increasing velocities (Fig 5(b)), indicating that biofilm morphology tends to be more flat in high-velocity fluid conditions as most of the mushroom protrusions can be removed.

**Fig 5.**
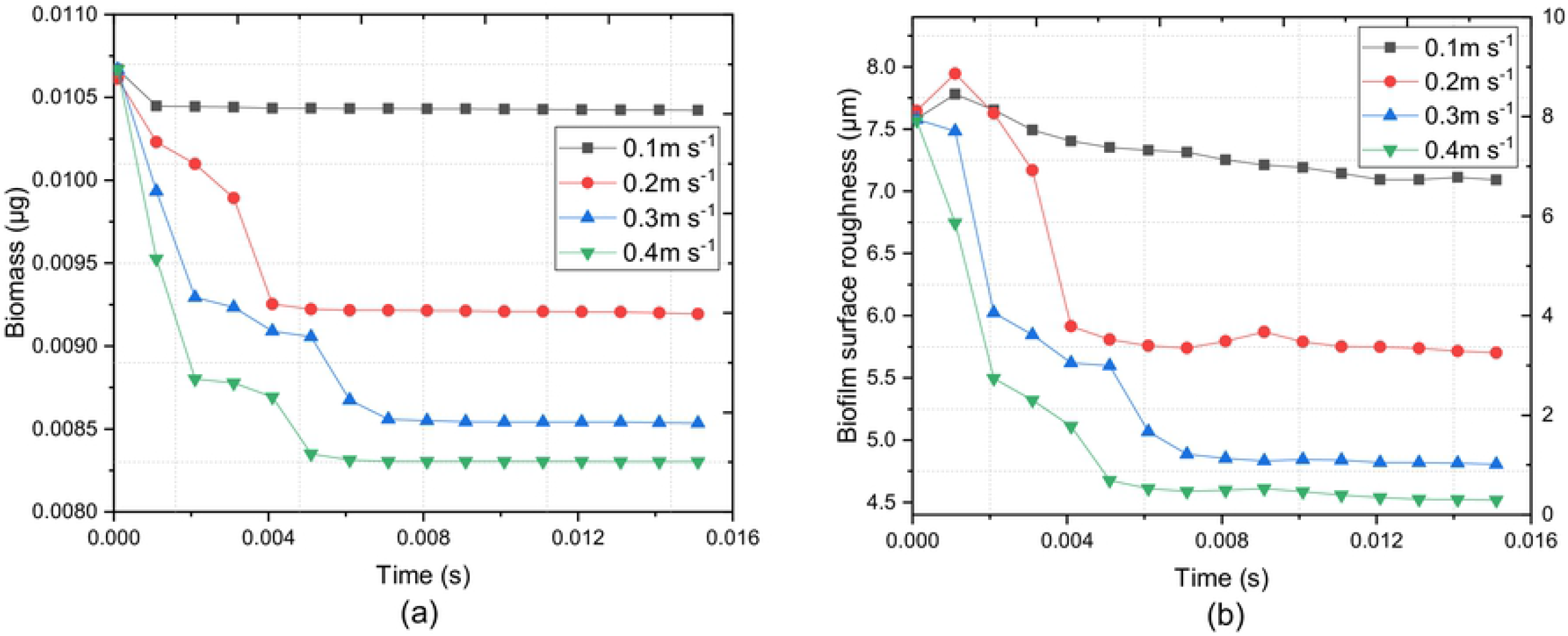
Effect of emergent properties on biofilm detachment: (a) total biomass, and (b) biofilm surface roughness.

### Case Study 2: Biofilm growth with 10^7^ particles

In this case study, we first show the development of a large and complex biofilm system and then focus on the parallel efficiency of the simulations. The aim of this case study is to demonstrate the capability and performance of NUFEB in the simulation of larger biological systems.

#### Biofilm development

The system is defined as a multi-functional group and multi-nutrient biofilm. In order to represent a more realistic biofilm, we explicitly consider nitrification as a two-step oxidation process that is performed by different groups of microbes: ammonia oxidizing bacteria (AOB) and nitrite oxidizing bacteria (NOB). In addition, the biofilm includes heterotrophs (HET) and their EPS production. The reaction model contains five soluble species, nutrients and products during microbial metabolism. The catabolic reactions include oxidation of ammonium NH_4_+ to nitrite NO_2_^−^ by AOB, oxidation of nitrite to nitrate NO_3_^−^ by NOB, and HET aerobic and anaerobic digestion by consuming organic substrate in oxygenated conditions or nitrate in anoxic denitrifying conditions. The kinetics and reaction stoichiometry of the modelled processes and their corresponding parameters are detailed in the S1 File. The computational domain is divided into three compartments. In the bulk compartment, nutrients are assumed to be completely mixed and their concentration is updated dynamically at each biological timestep, except for oxygen. We also assume that there is sufficient O_2_ and NH_4_^+^ but no NO_2_^−^ and NO_3_^−^ in the reactor influent. So the concentration of the two N compounds can only come from the transformation of NH_4_^+^. In the boundary layer compartment, a 20 *μm* distance from the maximum biofilm thickness to the bulk liquid is defined for solving the nutrient gradient. In the biofilm compartment, instead of using the super particle method [19], a small division diameter (1.3 *μm*) is chosen to represent real microbe sizes (on average 1 *μm* [44]).

The simulation is run on an in-house HPC system at Newcastle University. In Fig 6 and S5 Video we present the biofilm development over time. The system reaches 2.3 × 10^7^ particles after 160 hours (CPU time = 30 hours). The initial particles are randomly placed on the substratum. In the early stages of biofilm formation, due to high growth rate and sufficient supply of substrate from bulk liquid, heterotrophs grow faster than nitrifiers (0, 60 and 120 hours). As the biomass grows, the system turns to substrate-limited condition for the HET group, while there is still sufficient NH_4_+ due to its high initial concentration that favours nitrifier growth. As a result, biofilm surface coverage of heterotrophs becomes smaller than nitrifiers (160 hours). This phenomenon matches previous experimental results where the nitrifying population can be significantly higher than heterotrophs in a substrate-limited reactor [45]. The biofilm geometry forms a wavy structure after 160 hours. This is because of a self-enhancing process from the non-uniform initial microbial distribution [46]. The spatial distribution of NO_2_^−^ concentration is shown in Fig 7. It is clear that the NO_2_^−^ concentration field differs according to the AOB distribution, where regions with high NO_2_^−^ concentration are the locations where AOB clusters are present.

**Fig 6.**
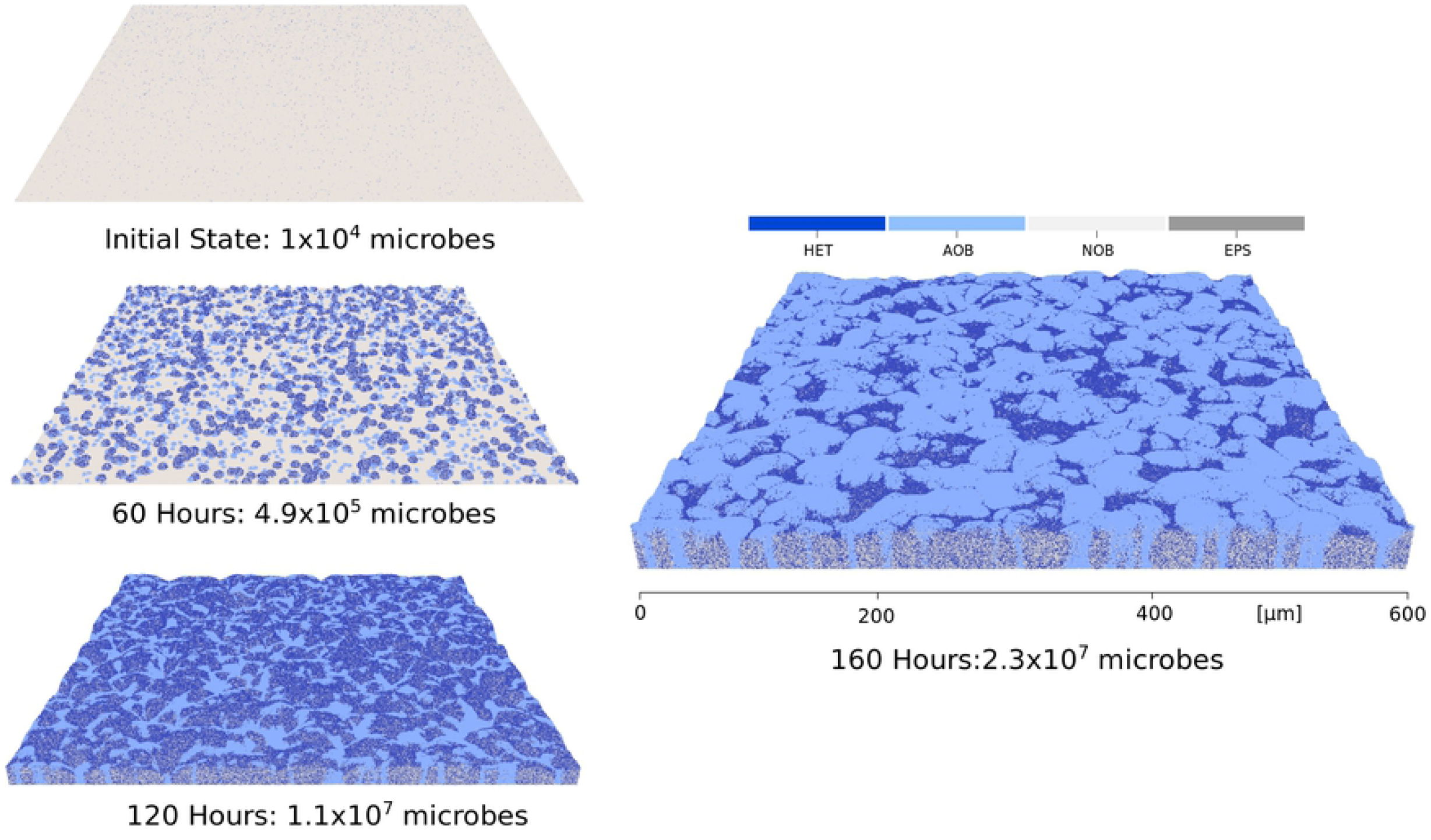
Biofilm development after 0, 60, 120, and 160 hours. The simulation uses 100 processors and 30 hours CPU time to reach 2.3 × 10^7^ particles. The biological timestep is 0.25 hour. Particle colours are blue for heterotrophs, grey for EPS, light blue for AOB, and light grey for NOB.

**Fig 7.**
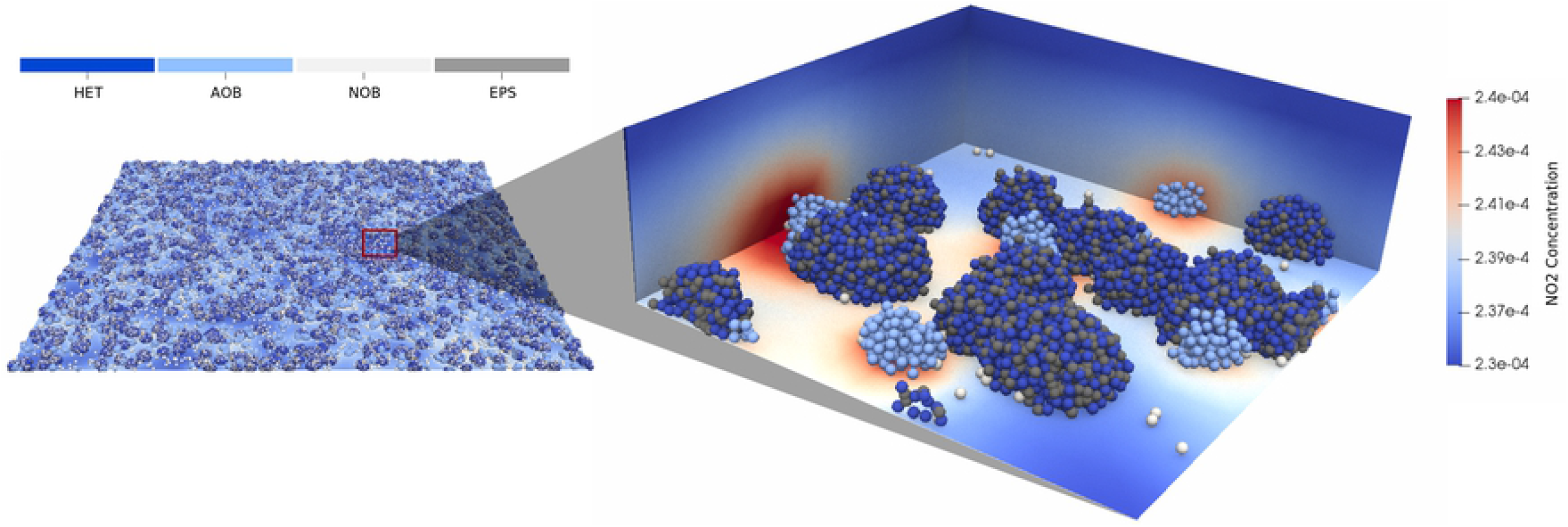
Nitrite concentration field at a small part of the simulated domain after 60 hours. The spatial distribution of NO_2_^−^ concentration follows the nitrifier distribution. The regions where NO_2_^−^ accumulates are due to production by AOB.

Fig 8(a) shows a quantitative evaluation of the total biomass accumulation over time. The trend shows linear biomass increase which indicates that the total microbial growth rate is not yet balanced by the decay rate. Therefore, a biomass steady state is not achieved after 160 hours. This is due to the high-oxygen environment (1 × 10^−2^ kg m^−3^) and the thin biofilm which nutrients can penetrate. However, the concentration of substrate and NH_4_^+^ in the bulk liquid relax to steady state before the total biomass concentration relaxes (Fig 8(b)), as bulk concentrations are mainly determined by biomass in the top biofilm layers, which ensures a high growth rate [19]. The NO_2_^−^ profile is influenced by both AOB synthesis and NOB consumption. Thus, the bulk concentration decreases with the increase of NOB populations.

**Fig 8.**
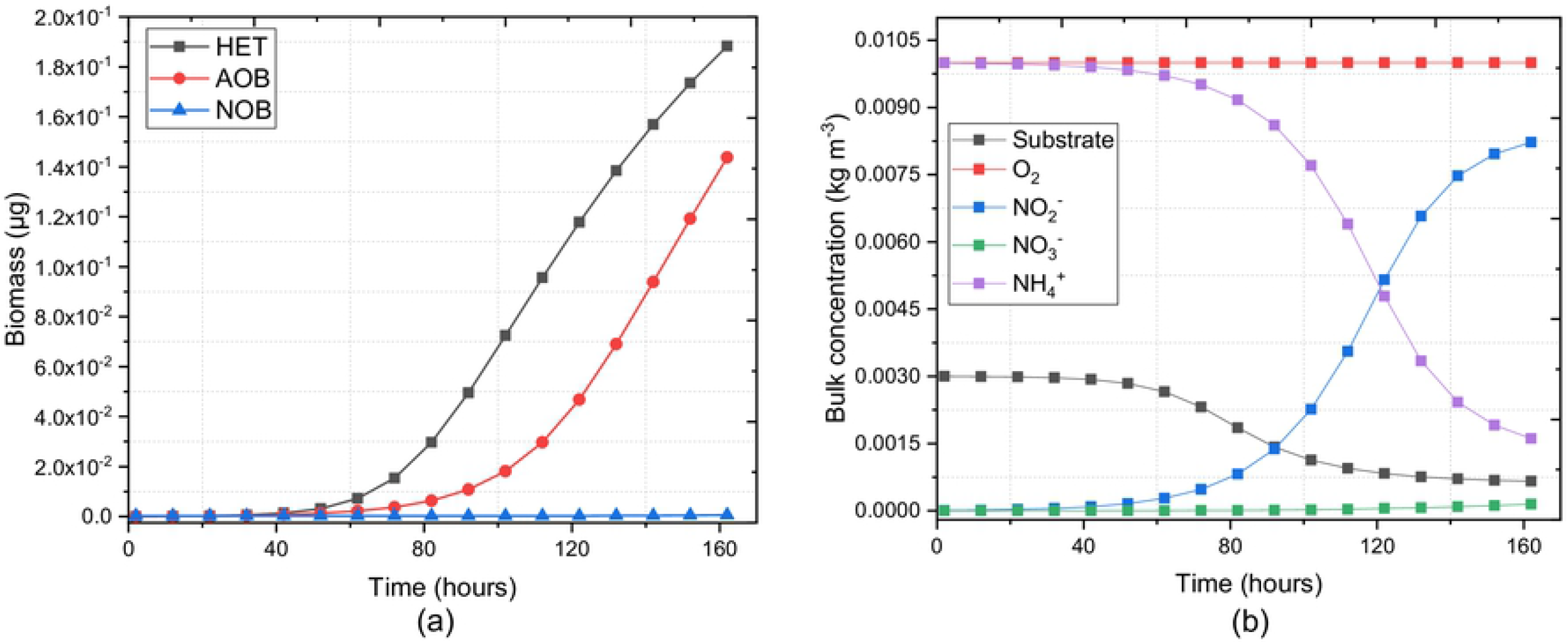
Quantitative evaluation of Case Study 2: (a) Total biomass of active functional groups over time, and (b) nutrients concentration in bulk liquid. Note that we assume the oxygen concentration in bulk liquid is kept constant by aeration.

#### Parallel performance

Parallel performance is crucial to IbM solvers for simulating large and complex problems. To investigate the performance of NUFEB, a scalability test was performed based on Case Study 2. The test aims at evaluating how simulation time varies with the number of processors for a fixed problem size. The *speed-up* of the scalability test is defined as *t_p0_/t_pn_*, where *t_p0_* and *t_pn_* are the actual CPU times spent on the baseline case and the test case respectively. Then the *parallel efficiency* is the ratio between the speed-up of the baseline and the test cases obtained when using a given number of processors, i.e., *N_p0_t_p0_*/*N_pn_t_pn_*, where *N_p0_* and *N_pn_* are the number of processors employed in these cases.

In order to reach a considerable number of microbes (2 millions), the baseline case is performed using 4 processors, and the subsequent tests were performed with increasing numbers of processors, ranging from 8 to 256. As mentioned previously, NUFEB implements two distinct solutions for the parallelisation of continuum-based and particle-based processes. The performance of the two solutions are studied separately, and are presented in Fig 9. It can be observed that when employing a small number of processors, both solutions are close to the ideal speed-up (linear increase) and the parallel efficiency is over 90%. As more processor nodes are added, each processing node spends more time doing inter-processor communication than useful processing, and the parallelisation becomes less efficient. In the tests using 128 and 256 processors, the parallel efficiency decreases to around 42%, but it still shows a good scalability.

**Fig 9.**
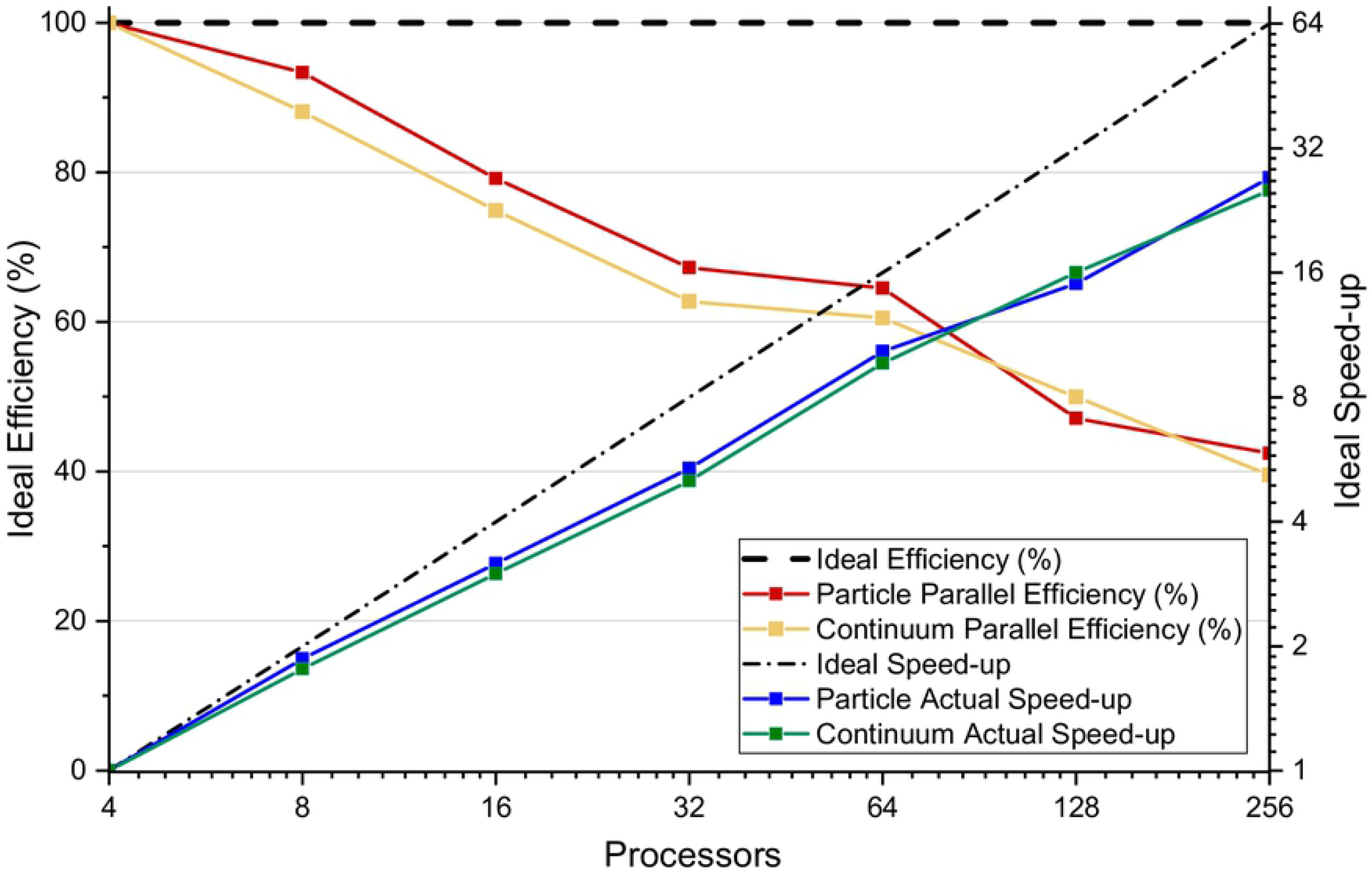
Performance of Case Study 2 with 4 – 256 processors. The initial conditions and the model parameters are kept the same in all cases. Parallel efficiency decreases with increasing numbers of processors due to inter-processor communication.

### Availability and future directions

In this paper, we have presented the NUFEB tool for modelling and simulating Individual Based models. The tool, documentation, and examples are publicly available on the GitHub repository: https://github.com/nufeb/NUFEB. To date, NUFEB has been adopted to model microbial communities in a variety of studies. In [17], we studied the influence of nutrient gradients on biofilm structure formation, and the influence of shear flow on growing biofilm. In [39] and [47], we used micro-scale NUFEB simulations to emulate the behaviour of microbial communities in the macro-scale. An emulation strategy for parameter calibration of NUFEB against iDynoMiCS has been applied in [48].

Ongoing work includes modelling and studying bacteria twitching motility, biofilm streamer oscillation and formation, and the influence of thermodynamics and pH on microbial growth. Our goal for NUFEB development in the future would be to deliver an even more general, efficient and user-friendly platform. This will include, for example, the development of an intuitive Graphical User Interface which will significantly improve the user experience. In order to make NUFEB available for power-efficient HPC architecture, we will also focus on a Kokkos port for the NUFEB code. This would allow the code to run on different kinds of hardware, such as GPUs (Graphics Processing Units), Intel Xeon Phis, or many-core CPUs.

The computational demands of IbM will always place a limit on the scale at which they can be applied. However, it is now evident that this limit can overcome by the use of statistical emulators [39]. A statistical emulator is a computationally efficient mimic of an IBM that can run thousands of times faster. In principle this new approach will allow the output of an IBM to be used at the metre scale and beyond and thus to make predictions about systems level performance in, for example, wastewater treatment or biofilm-fouling and drag on ships. This is a strategically important advance that creates a new impetus for the development of IbM that can credibly combine chemistry, mechanics, biology and hydrodynamics in a computationally efficient framework. NUFEB is, we believe, the first generation of IbM to meet that need and will help IbM transition from a research to application.

## Supporting information

**S1 File. Supporting information (SI).**

**S1 Manual. User manual.**

**S1 Video. Growth of a biofilm without flow**. The biofilm is initially grown for 9 days without flow. It forms a mushroom-shaped structure in an oxygen-limited condition (1 × 10^−4^ kg m^−3^).

**S2 Video. Biofilm removal at U_*f*_ = 0.2 m s^−1^**. Large chunks of microbes detach from the biofilm surface and then remove from the systems. The biofilm morphology changes from a rough to a flat surface.

**S3 Video. Biofilm removal at U_*f*_ = 0.1 m s^−1^**. The top of the biofilm is highly elongated. Small clusters erode from the deforming biofilm and the amount of biomass removed is less than the **U**_*f*_ = 0.2 m s^−1^ case.

**S4 Video. Biofilm deformation and detachment with periodic wall at U_*f*_ = 0.2 m s^−1^**. Microbes crossing the boundary will re-appear on the opposite side of the domain. It can be observed that the detached clusters can re-attached to the biofilm surface or re-agglomerate with other clusters.

**S5 Video. Growth of a large biofilm system**. The multiple functional groups biofilm is grown without flow and reaches 2.3 × 10^7^ particles after 160 hours (CPU time = 30 hours). The biofilm forms a wavy structure because of the high nutrient concentration environment and the non-uniform initial microbial distribution.

## Acknowledgments

The work is supported by the UK’s EPSRC EP/K039083/1 Newcastle University Frontiers in Engineering Biology (NUFEB) project. The authors would like to thank Dr. Prashant Gupta and Dr. Curtis Madsen for the initial implementation of NUFEB. We also thank the NUFEB modelling team and Yuqing Xia for their helpful advice.

## Footnotes

1 If energy-growth model is applied, the computation is accomplished with pH dynamics and gas-liquid transfer to update the concentration of dissociation forms, pH and reaction rate.

